# New insights on the genetics of hair whorls from twins and the Southern hemisphere

**DOI:** 10.1101/2023.02.20.529302

**Authors:** Marjolaine Willems, Quentin Hennocq, Juan José Cortés Santander, Roman Hossein Khonsari

## Abstract

The mechanisms determining the rotation direction and position of hair whorls are unknown. Here we report observations on twins suggesting that the morphological parameters of whorls have genetic bases, and provide comparative data on whorls from children born in the Northern and Southern hemispheres, indicating that whorl formation also depends on environmental factors. Our results underline the importance of unusual morphological phenomena for providing general information on normal developmental processes, and plead for large-scale epidemiological assessments to support our surprizing initial findings.

---

According to the hedgehog theorem of algebraic topology (Renteln 2013), most hairy animals have hair whorls: on full spheres, you cannot comb a hairy ball flat without creating a cowlick. In humans, hair whorls are constant on the scalp, even though this surface is a spherical cap rather than a full sphere. Whorls can be unique or multiple, are generally lateralized, and oriented either clockwise or counterclockwise. Their developmental origins and the mechanisms determining their rotation direction are unclear. Both genetics and physical constraints could be implicated in determining the location and orientation of whorls, as in other lateralisation phenomena (Forrest et al. 2022). As is Blaschko lines, which include whorl-like distributions on the scalp surface (Happle 2013), hair whorls may be related with cell migration processes but there is no current experimental evidence for this hypothesis. A report based on 404 new-borns showed that 98.5% of babies had single parietal whorls and 1.5% had double whorls, with 93.8% of clockwise whorls (Wunderlich & Heerema 1975). The most common configurations were, listed by decreasing prevalence: clockwise central, clockwise right, clockwise left, counterclockwise central, double, counterclockwise left, and counterclockwise right. Whorls have been associated with a surprising list of physical and cognitive characteristics: behaviour in cattle (Górecka et al. 2007), laterality in humans (Klar 2003) and even homosexuality (Klar 2004) but these reports have been invalidated (Rahman et al. 2009; Çetkin et al. 2020). Several clinical arguments indicate that whorl formation may be at least partially under genetic control. For instance, atypical whorls (double or multiple, frontal hair whorls) seem to be more prevalent in patients with genetic conditions (de Leeuw et al. 2010). Twinning is a potential model to investigate the genetic bases of whorl distribution, by comparing monozygotic and dizygotic cases, even though monozygotic twins arise both from monochorionic pregnancies and from bichorionic pregnancies, with distinct mechanisms that could potentially interfere with whorl formation. Interestingly, whorls are regularly mentioned as a physical feature in mirror twins – 25% of monozygotic pairs – without supporting data in the literature. Furthermore, double occipital hair whorls seem to be more prevalent in twins than in singletons, and may be specifically prevalent in monozygotic twins (Paine et al. 2004).

In order to investigate the developmental origins of whorls, we performed an observational study in twins from monochorionic pregnancies (which are 100% monozygotic), and twins from bichorionic pregnancies of the same sex (which are 20% monozygotic and 80 % dizygotic) (Yeaton-Massey et al. 2021; Hall 2003). In order to evaluate environmental effects on whorl rotation direction, we compared whorl configuration in a population born in the Northern hemisphere (Paris, France) with a population born in the Southern hemisphere (Santiago, Chile). We obtained unexpected results.

Three populations were considered: (1) Northern hemisphere general population (50 children [25 girls and 25 boys], aged 1-10, born in Paris, France, admitted for craniofacial trauma at Necker – Enfants Malades University Hospital (Paris, France), without medical history, from March 1st to March 31st 2021); (2) Southern hemisphere general population (similar inclusion criteria, children born in Santiago, Chile, admitted in Clinica Universitad de los Andes (Santiago, Chile)); and (3) same-sex Northern hemisphere twins (all pairs of same-sex twins born in Necker – Enfants Malades University Hospital (Paris, France) from January 1st 2022 to March 31st 2022). Whorl rotation direction (clockwise, counterclockwise) and whorl position (left, right, central) were recorded, as well as twinning type: BiChorial BiAmniotic (BCBA), MonoChorial BiAmniotic (MCBA), and MonoChorial MonoAmniotic (MCMA). Univariate logistic models were designed for each explanatory variable to screen for a statistical association with rotation direction and position. For twins, the variable of interest was binary, i.e. same rotation direction (reference class) or opposite directions for each twin pair. For controls, all single children combinations were included as virtual twins. New odds ratios (OR) were calculated and compared for both hemispheres. The significance threshold was p < 0.05; a significant parameter influenced the relevant variables for each model. Assumptions of normality and homoscedasticity of errors were tested. The statistical analyses were performed on R 3.6.21 using the *nlme2* and *ggplot3* packages.

Seventy-four (37 pairs) twins were included, and absolute ratio values seemed to indicate that most whorls were clockwise and rotated in the same direction within pairs (Table 1). The OR for opposite rotation directions between two twins was ≠1 (p=0.017), meaning that whorls rotated preferentially in the same direction in twins. The OR for different whorl laterality between two twins was not ≠1 (p=0.869), meaning that whorls lateralized either on the same side or on opposite sides in twins. Sex, pregnancy type, number of chorions and whorl position did not affect rotation direction (p<0.05) and position (p<0.05). ORs were <1 for Northern and Southern hemispheres, meaning that whorls rotated preferentially in the same direction in simulated twins. Of note, OR=0.04 [0.03; 0.05] for simulated twins in the Northern hemisphere, less than OR for real twins (0.42 [0.20; 0.83], p < 0.001) with no confidence interval superimposition: it was more likeable to find opposite whorls in twins than in a random pair of children (p < 0.001). Also, OR for the Northern hemisphere (0.04 [0.03; 0.05]) was less than the OR for the Southern hemisphere (0.28 [0.24; 0.32]) with no confidence interval superimposition, which meant than counterclockwise whorls were more frequent in the Southern hemisphere (p < 0.001).

**Table 1.**
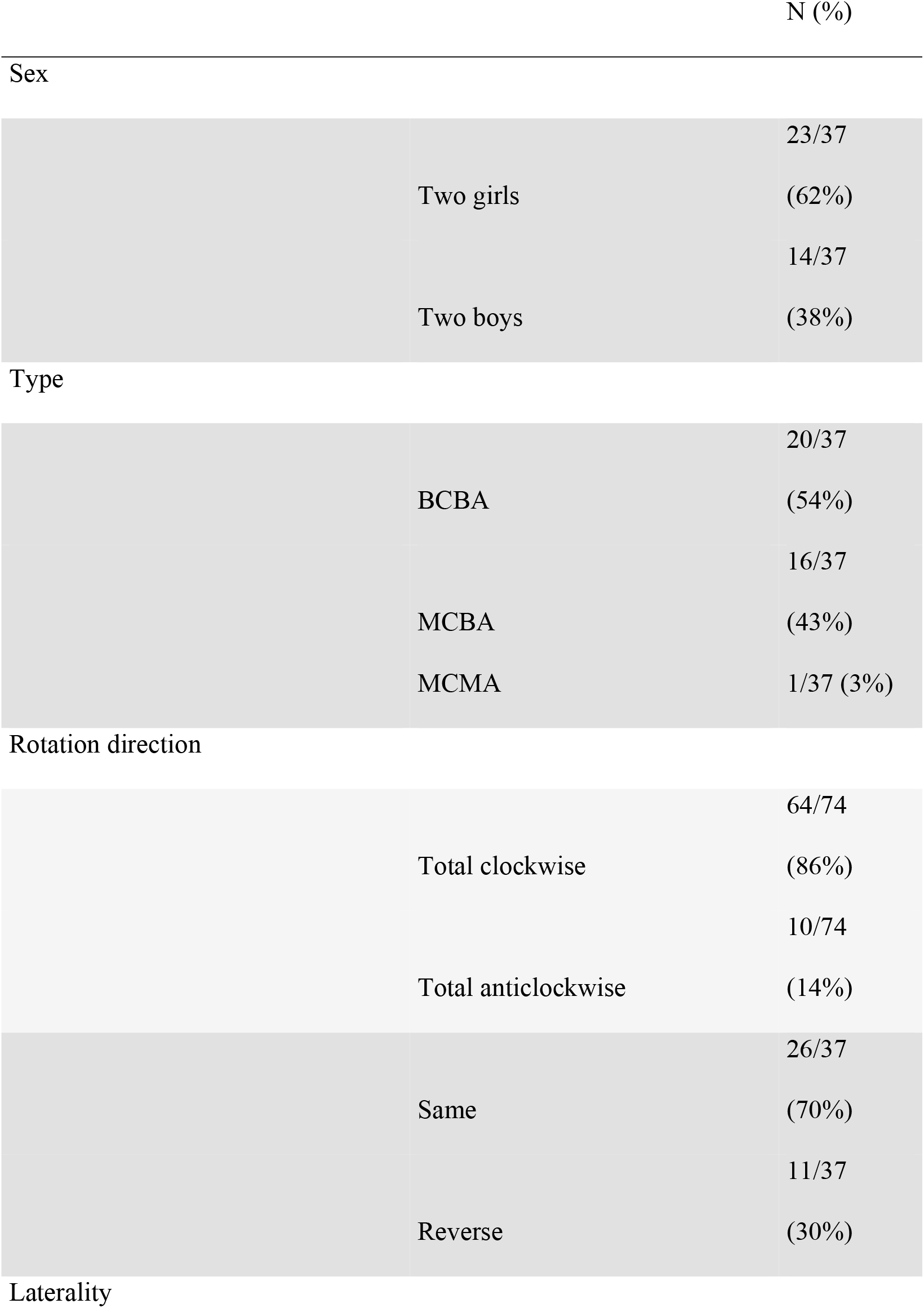
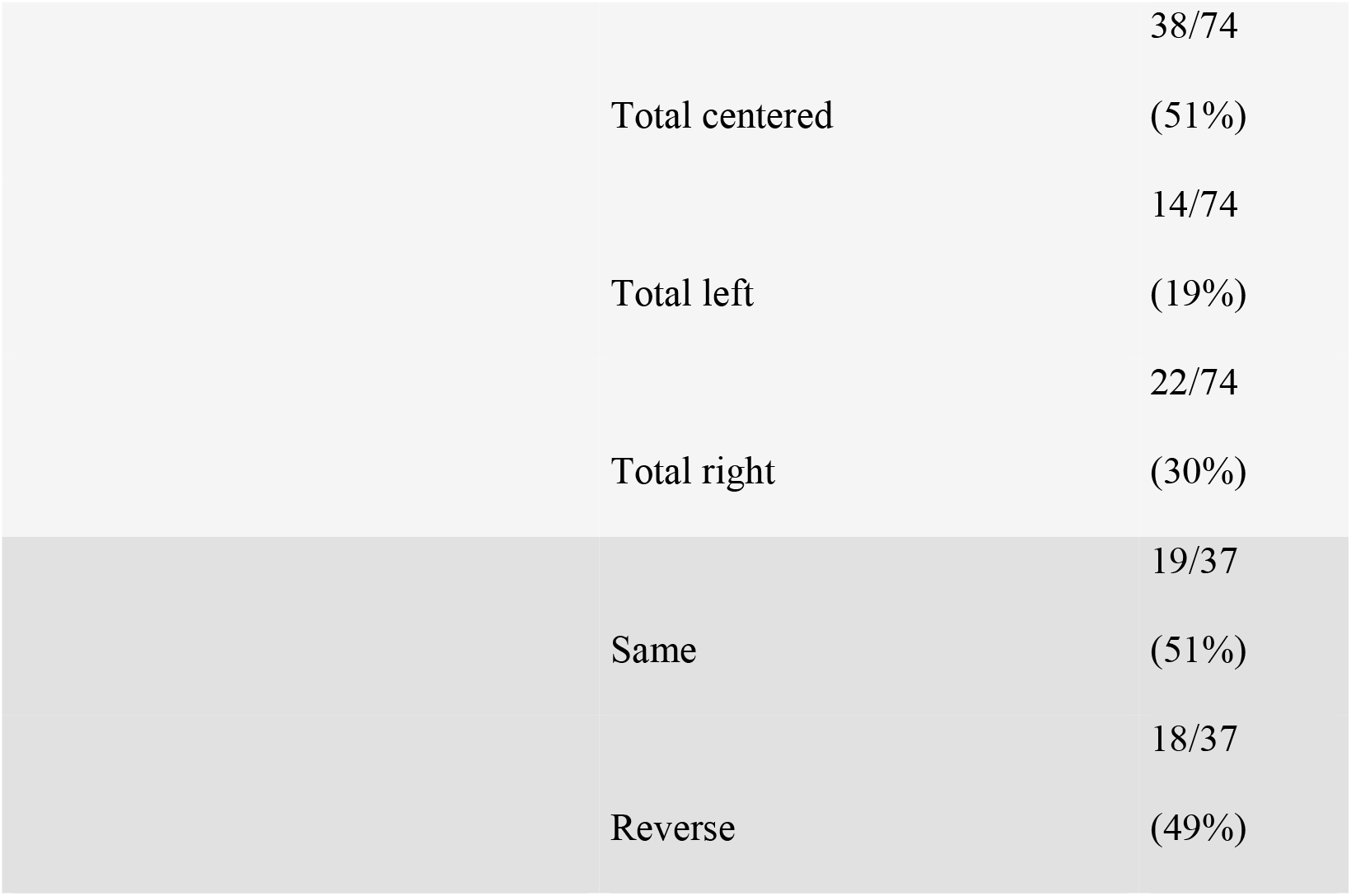
Data description. The denominator is 74 to denote the total number of individuals and 37 to denote the total number of pairs of twins. BCBA = Bichorial Biamniotic; MCBA = Monochorial Biamniotic; Monochorial Monoamniotic.

**Table 2.**
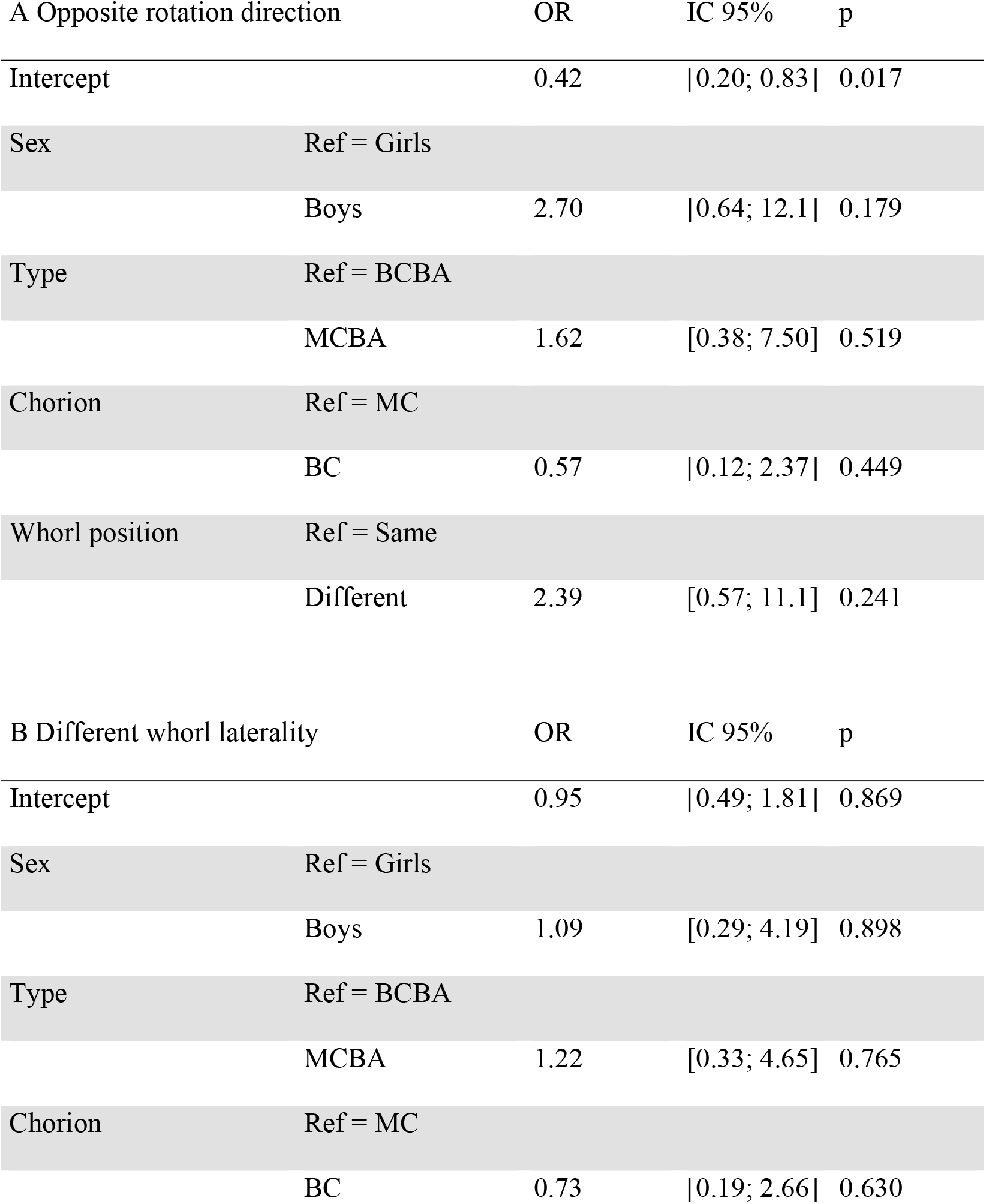
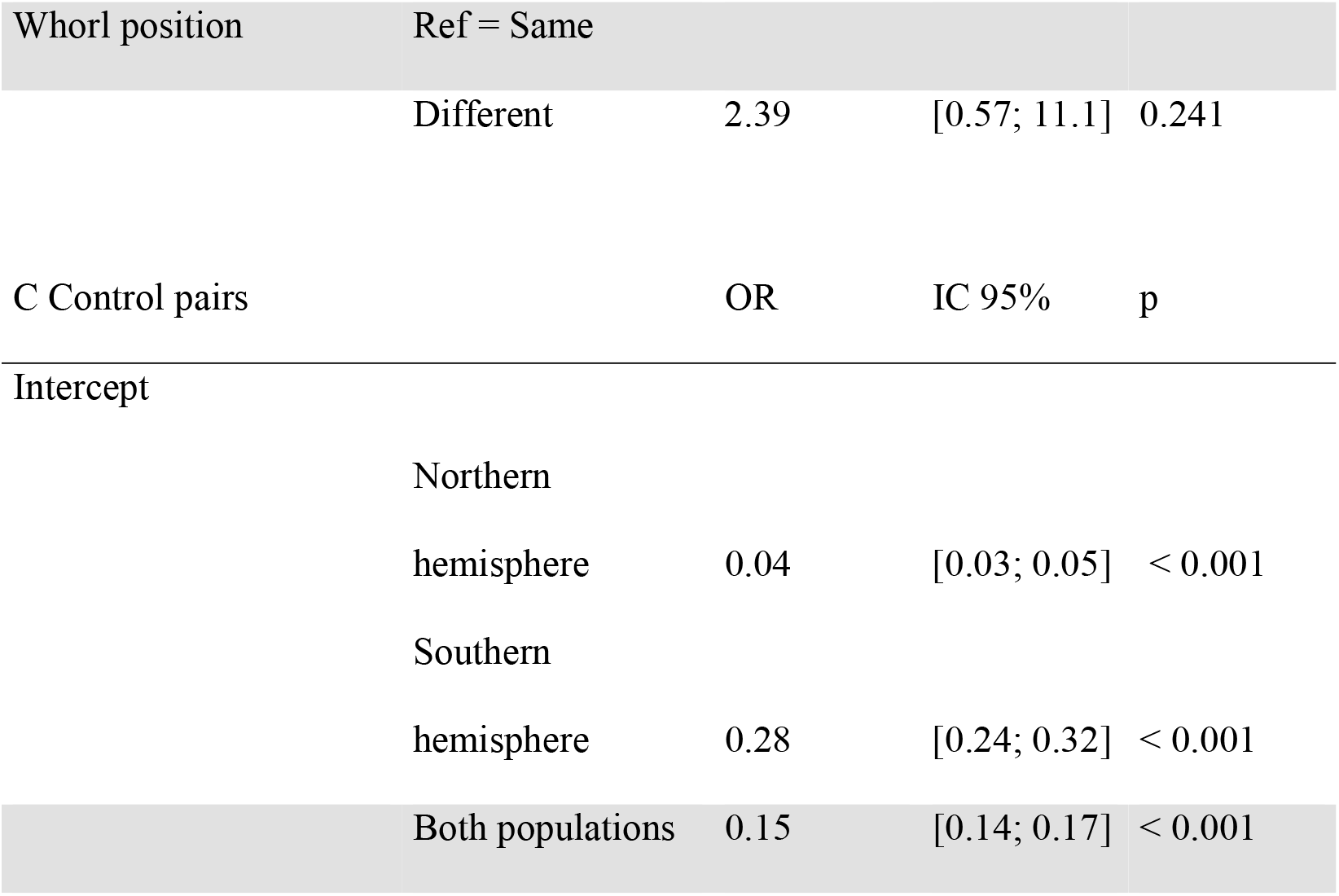
A. Rotation direction in twins. Intercept value corresponded to the OR of opposite rotation directions between two twins. B. Laterality in twins. Intercept value corresponded to the OR of different whorl laterality between two twins. C. Control pairs. BCBA = Bichorial Biamniotic; MCBA = Monochorial Biamniotic; Monochorial Monoamniotic, BC = Bichorial; MC = monochorial.

Preferential rotation direction of hair whorls in twins from monochorionic pregnancies and same-sex bichorionic pregnancies indicated that whorl characteristics were partially determined by genetic factors. A higher prevalence of counterclockwise whorls in the Southern hemisphere underlined the potential role of environmental factors, such as external physical constraints. It was not clear whether this hemispheric effect on rotation direction could be related to the Coriolis force. Classical experiments on liquid drainage show counterclockwise rotation in the Northern hemisphere (Shapiro 1962) and clockwise rotation in the Southern hemisphere (Trefethen et al. 1965). As multiple physical forces are involved in the initial steps of craniofacial development (Fleury 2022), there is probably no straightforward link between the rotation direction of whorls and any type of physical phenomenon potentially influenced by the Coriolis force. Nevertheless, this hemispheric effect questions the nature of mechanical forces involved in early development.

This report should lead to furthers investigations on the origins of hair whorls and more generally on the physical and chemical mechanisms determining the three-dimensional distribution of cell populations during development. Similarly, other enigmatic patterns such as concentric rings (Baló’s concentric sclerosis, concentric skin rings in mycoses) or annular ‘wood-grain’ lesions (*erythema gyratum repens*) could provide critical information on general developmental processes by providing unique insights on phase transitions, critical points and the factors influencing order parameters. Large-scale epidemiological assessments in genetically confirmed monozygotic twins and on both hemispheres would be interesting to confirm our initial results and better understand the interactions between environmental factors and early developmental mechanisms.

## Acknowledgements

Many thanks to Laurine Aljancic, Eléonore Bobo, Florent Fuchs, Justine Giunta, Marine Huby, Nicolas Kogane, Camille Larrieu-Arguillé, Thibaud Mernier, Tan Mai Nguyen, Romy Razssiguier, Roberto Requena, Valentin Ruault, Eleonora Segna, Julien Stirnemann, Vincent Sounthakith, and Sara Tunon de Lara for data collection. Many thanks to Eve, Thaïs, Roxane and Cléophée for having inspired this study, and to all patients for their contribution.

